## Supplementary Text for "Surface frustration re-patterning underlies the structural landscape and evolvability of fungal orphan candidate effectors"

1. **Analysis of the OCE structural network**

The structural similarity network including 2 561 orphan effectors with at least three analogs at Z≥5.2 featured nodes of high connectivity (**Fig S1A**), corresponding to protein folds shared between multiple orphan effectors. Six effectors from the network had experimentally determined structures (**Fig S1B**): *Blumeria graminis* ribonuclease (RNase)-like fold (RALPH) CSEP0064/BEC1054 (6fmb) [1], *Trichoderma* Tsp1 (7esw) [2], *Zymoseptoria tritici* KP6 killer toxin (6qpk) [3], *Drechmeria coniospora* g7941 (6zpp), and *Pyricularia oryzae* Avr-Pik avirulence effector (6fud) [4] and Hrip2 elicitor (5fid) [5]. The match between experimentally determined structures and AlphaFold2 predictions was very good with RMSD ranging from 0.468 and 3.94 angstroms. We conclude that OCEs group into a few highly abundant folds at the Kingdom level, with multiple OCEs being close structurally in spite of reduced sequence homology.

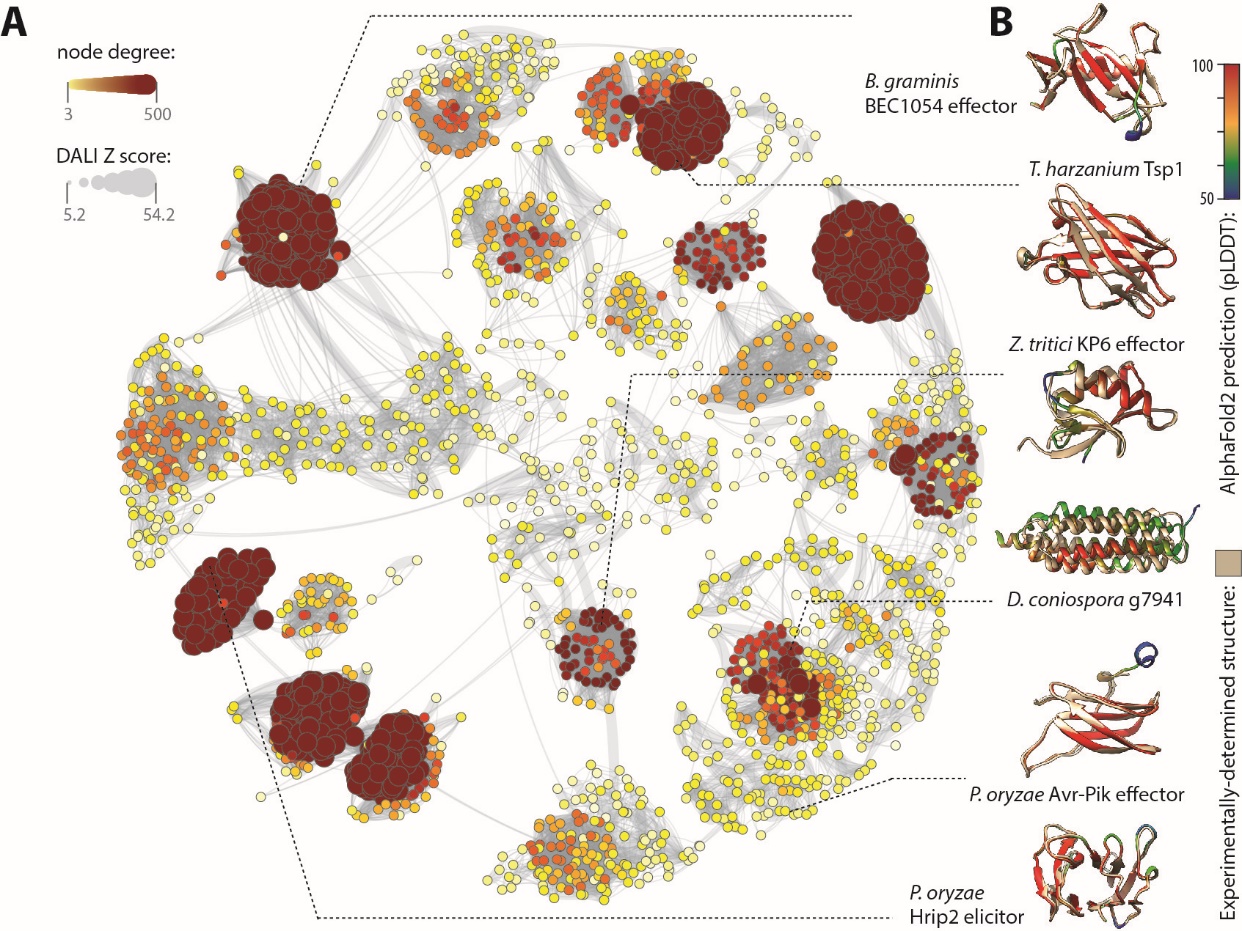

**Figure S1. Structural relationship between orphan candidate effectors in 20 fungal genomes.**  **(A)** The structural landscape of fungal orphan candidate effectors shown as a similarity network. Vertices are OCE predicted structures colored and sized according to their number of edges which correspond to pairwise Dali Z-scores. **(B)** OCEs of the network for which an experimentally-determined structure is available. Predicted structures colored according to pLDDT score are superimposed with experimentally determined structures (tan color). Dotted lines point towards the corresponding vertices in the network.

Most of the broadly conserved fold covered the full length of OCEs. Exceptions included the short helical folds (1-helix and 3-helix) which were associated with other structural features in some OCEs, reducing the overall structural similarity for proteins harboring these folds (**Fig S1A**).

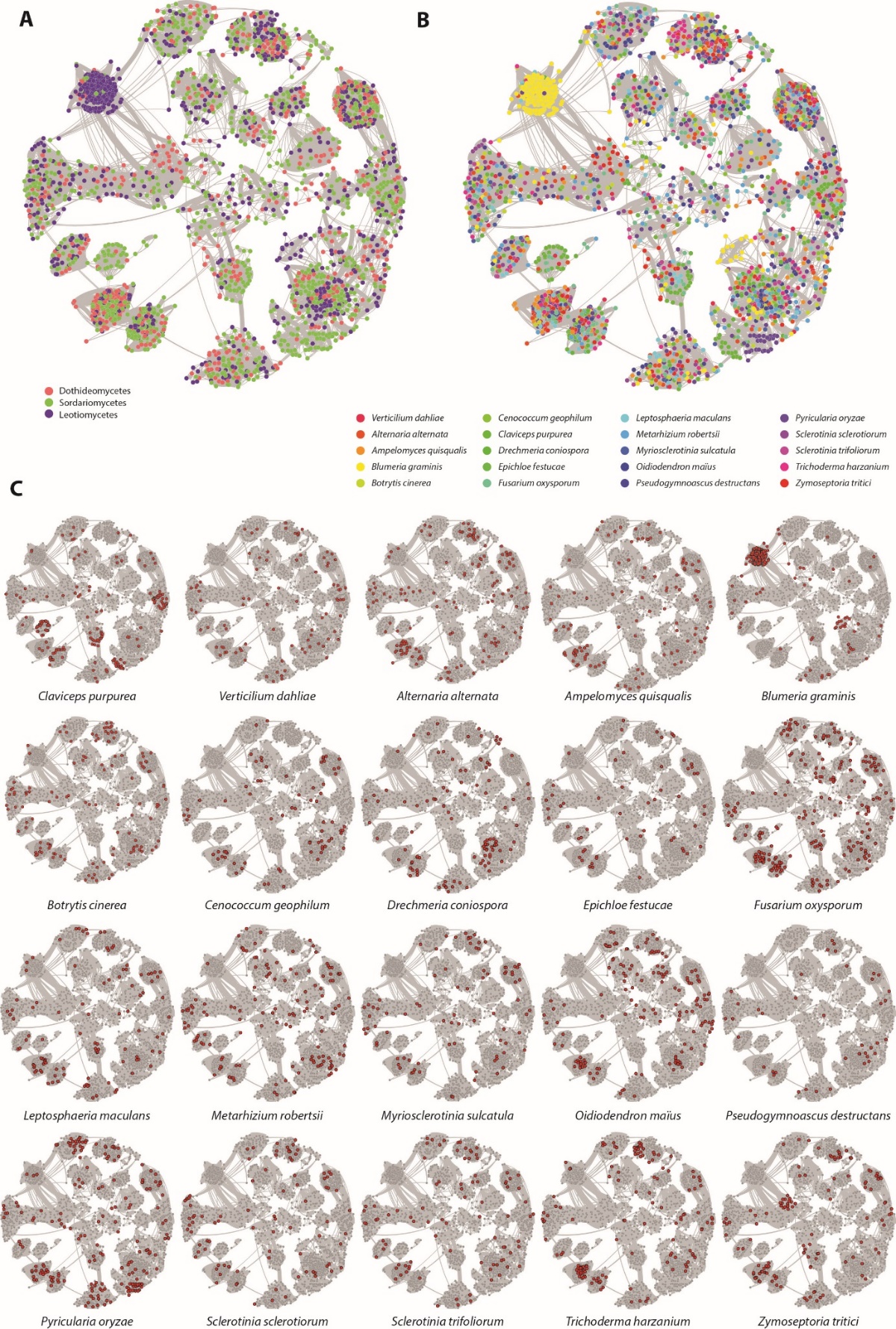

**Figure S2. Lineage and species representation across the OCE structural similarity network. (A)** Mapping of the three major fungal lineages onto the structural similarity network, with nodes colored according to the lineage they belong to. **(B)** Mapping of the twenty fungal species onto the structural similarity network, with nodes colored according to the species they belong to. **(C)** Structural similarity network with nodes corresponding to each of the twenty species highlighted.

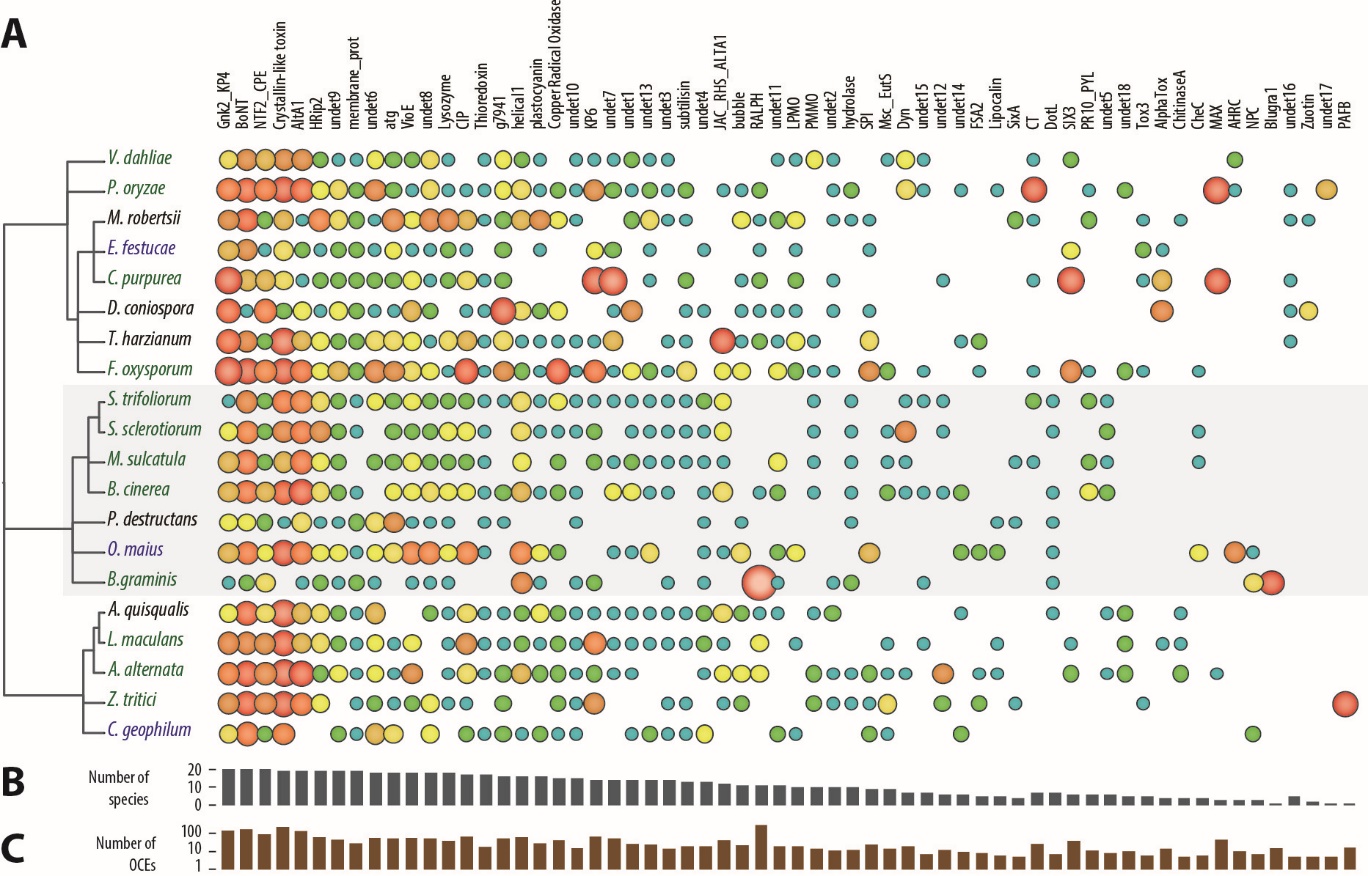

**Figure S3. Species distribution of the 62 major OCE structural groups. (A)** Number of OCEs from each of the 62 major family in the 20 fungal species analyzed (see also Table S4). Number of occurrences ranges from 1 (light blue) to 262 (RALPH in *B. graminis*, red). **(B)** Number of species containing at least one member of the OCE family. **(C)** total number of OCEs in each family.

Out of 62 major folds, only three were specific to a single fungal species (**Fig. S2, S3**), suggesting that specialization in these OCEs is limited. Fourty four OCE folds were detected in Leotiomycetes, Sordariomycetes and Dothideomycetes, with only five OCE folds were restricted to a single lineage. Folds enriched in the Sordariomycetes included MAX, Alpha Toxin and Tox3, folds enriched in the Leotiomycetes were RALPH, Blugra1, and DotL, and folds enriched in the Dothideomycetes were PAFB and ChitinaseA. Thirty-five OCE folds were detected in pathogens infecting plant, fungal and animal hosts, while 13 OCE folds were restricted to fungi infecting only one of these host groups. No OCE fold was specific to mycoparasites, the Zuotin-like fold was the only one restricted to animal pathogens, 12 folds were restricted to plant pathogens including MAX and SIX3. Fourty-six OCE folds were detected in pathogens with a mutualistic, a necrotrophic and other pathogenic lifestyles. With the exception of the KP6 fold absent from animal pathogens, all the seventeen most abundant OCE folds were detected in fungi from all lineages, infecting all host groups and with all lifestyles. Expression profiling of *S. sclerotiorum* OCEs in diverse *in vitro* and *in planta* conditions (**Fig. S4**) shows that several OCEs are preferentially expressed during plant infection, some are expressed preferentially *in vitro*, others are equally well expressed across all conditions tested.

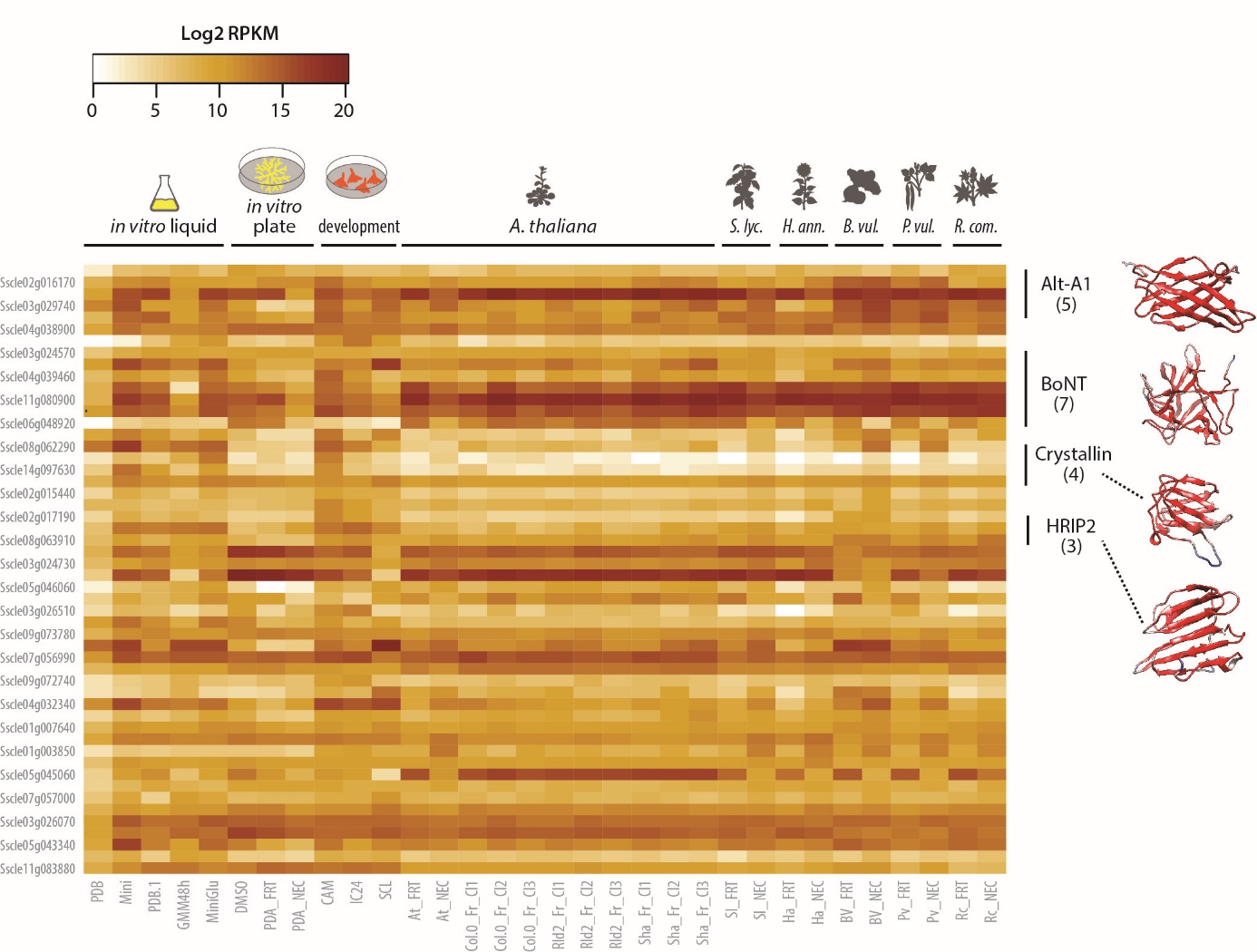

**Figure S4. Expression of genes encoding OCEs in *Sclerotinia sclerotiorum* in 11 *in vitro* and 21 *in planta* conditions**. RPKM, reads per kilobase per million; *A. thaliana*, *Arabidopsis thaliana*; *S. lyc*., *Solanum lycopersicum*; *H. ann*., *Helianthus annuus*; *B. vul*., *Beta vulgaris*; *P. vul*., *Phaseolus vulgaris*; *R. com*., *Ricinus communis*.

2. **Estimate of the number of OCE folds**

In order to estimate the total number of OCE folds in the fungal kingdom, we randomly sampled 1 to 19 fungal genomes and counted the number of OCE folds detected (**Fig S5**). We then used a logarithmic regression on the number of OCE folds detected according to the number of fungal genome sampled. Assuming our sampling of fungal species is representative of the actual diversity, we would expect a maximum of 70 OCE folds to exist across 2 million fungal species.

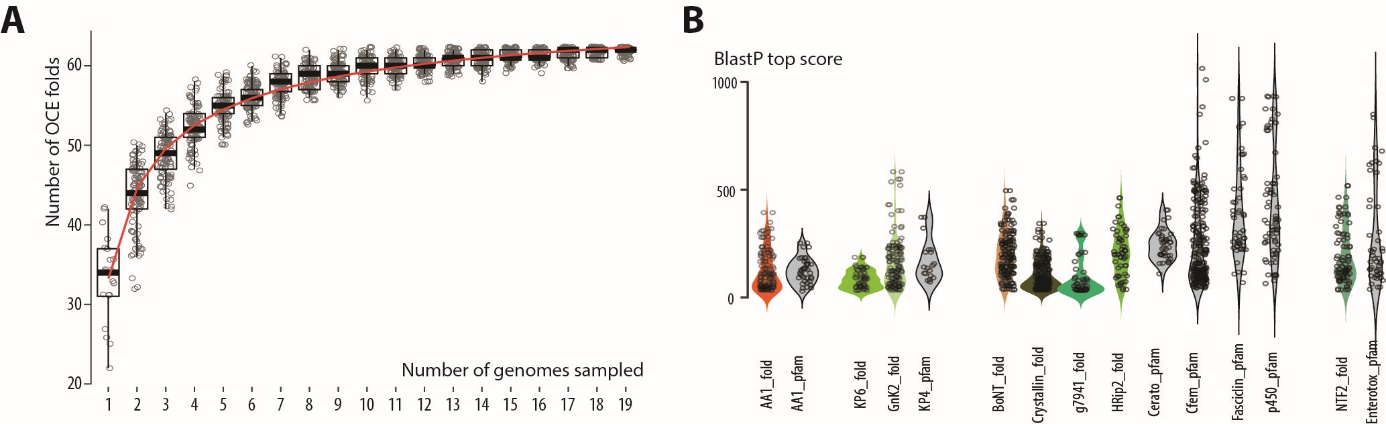

**Figure S5. Logarithmic regression on the number of distinct OCE folds detected according to the number of fungal genomes sampled.** The red line shows logarithmic regression. Dots show the number of distinct OCEs (out of the 62 from the complete dataset) in 100 random samples of the 20 fungal genomes. Boxplots show median values (thick line), first and third quartile values (box) and 1.5 times the interquartile range (whiskers).

3. **Sequence-structure similarity relationship**

OCEs adopting a same conserved fold often showed limited protein sequence identity. In the AA1/PEvD1/TSP1 fold for instance, the average pairwise Z-score was 22.9 while the average amino-acid identity was 10.1%. To characterize the relationship between sequence similarity and structure similarity in OCEs, we first compared the sequence similarity of OCE closest homologs within the 20 proteomes with that of functionally-related proteins sharing a common PFAM domain (**Fig. S6**). Overall, top BlastP score for OCEs tended to be skewed towards lower values than PFAMs, but a few OCEs also showed close homologs.

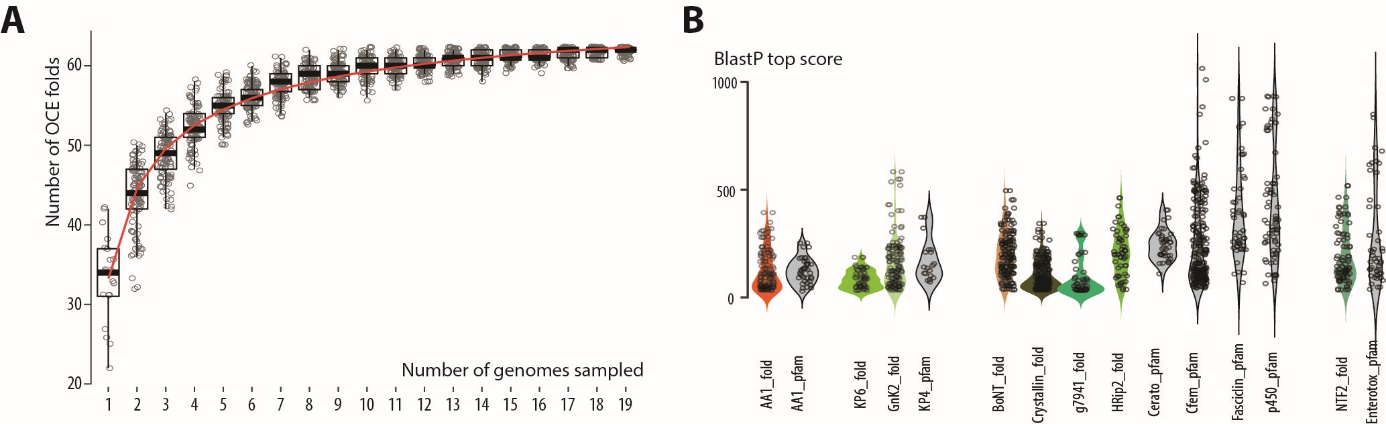

**Figure S6. Comparative analysis of similarity assessed by BlastP between OCE families and selected PFAMs from the 20 fungal genomes.** Data points show the top non-self BlastP score against the combined proteome of the 20 fungal species. Entries with the ‘_fold’ suffix correspond to OCE families, the ‘_pfam’ suffix denotes proteins sharing a same PFAM domain.

Next, we assessed the relationship between sequence similarity and structural similarity in seven OCE families and three control protein families (**Fig. S7**). We focused on pairwise protein comparisons above the 97.5% prediction interval as remarkable for highlighting proteins with high structural similarity in spite of significant sequence divergence.

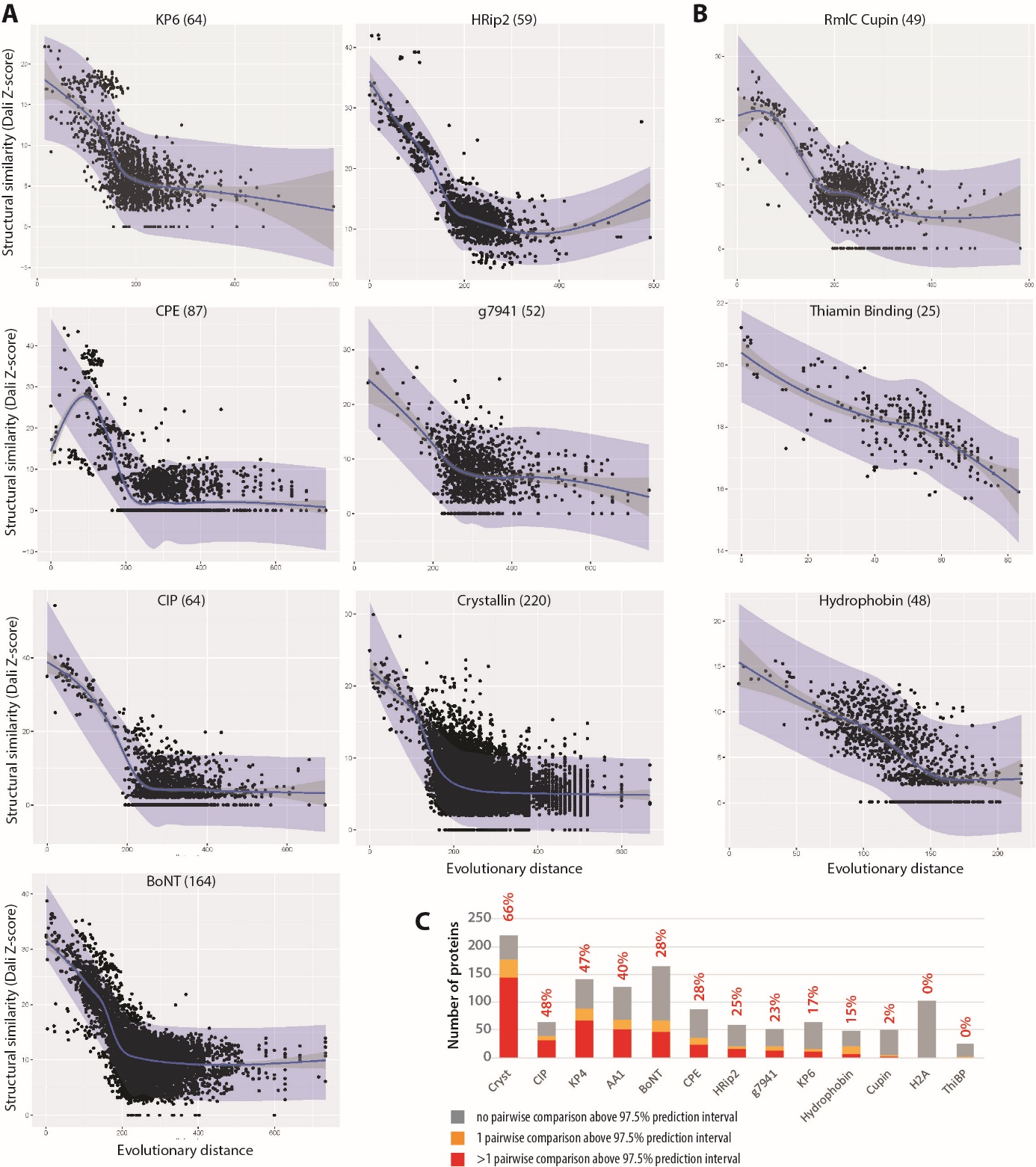

**Figure S7. Relationship between structural similarity and sequence similarity in OCEs.** Structural similarity (Dali Z-score) according to evolutionary distance (Jukes-Cantor corrected distance calculated on curated sequence alignments) for seve OCE groups **(A)** and three housekeeping control groups **(B)**. The green dotted lines is the local average. Number in brackets indicate the number of protein structures compared. (C) Number of proteins with 0, 1 or more than 1 pairwise comparison above the 97.5% prediction interval. The percentage of proteins with 1 or more comparison above the 97.5% prediction interval is indicated above each bar.

4. **Profile HMM-HMM comparisons demonstrate deep ancestry of most members of OCE structural families**

Several studies have shown that amino acid sequences evolve faster than protein structures. Highly divergent structurally similar proteins with common ancestors may not be detected using standard methods such as BLAST. To assess whether divergent OCEs structurally similar to one another exhibited distant common ancestry we used sensitive profile hidden Markov model (HMM) comparisons. For this analysis, we used BLASTp to identify up to 1,000 homologues from the NCBI nr database fungal division of each of the OCEs across 12 of the largest OCE clusters, including AltA1, atg, BoNT, CIP, Crystallin, g7941, Gink2 / KP4, HRip2, KP6, NTF2 / CPE, SIX3 and VioE.

We identified 593,597 BLASTp hits to these OCE families and grouped them into clusters with >= 80 % coverage of all members and a similarity of >= 30 %. This generated 3,467 clusters made up of 590,032 proteins and 3,565 singletons that did not meet these alignment criteria. We then conducted all vs all pairwise HMM-HMM comparisons between clusters and sequence-HMM comparisons between singletons and clusters for each of the families (**Fig. S8**).

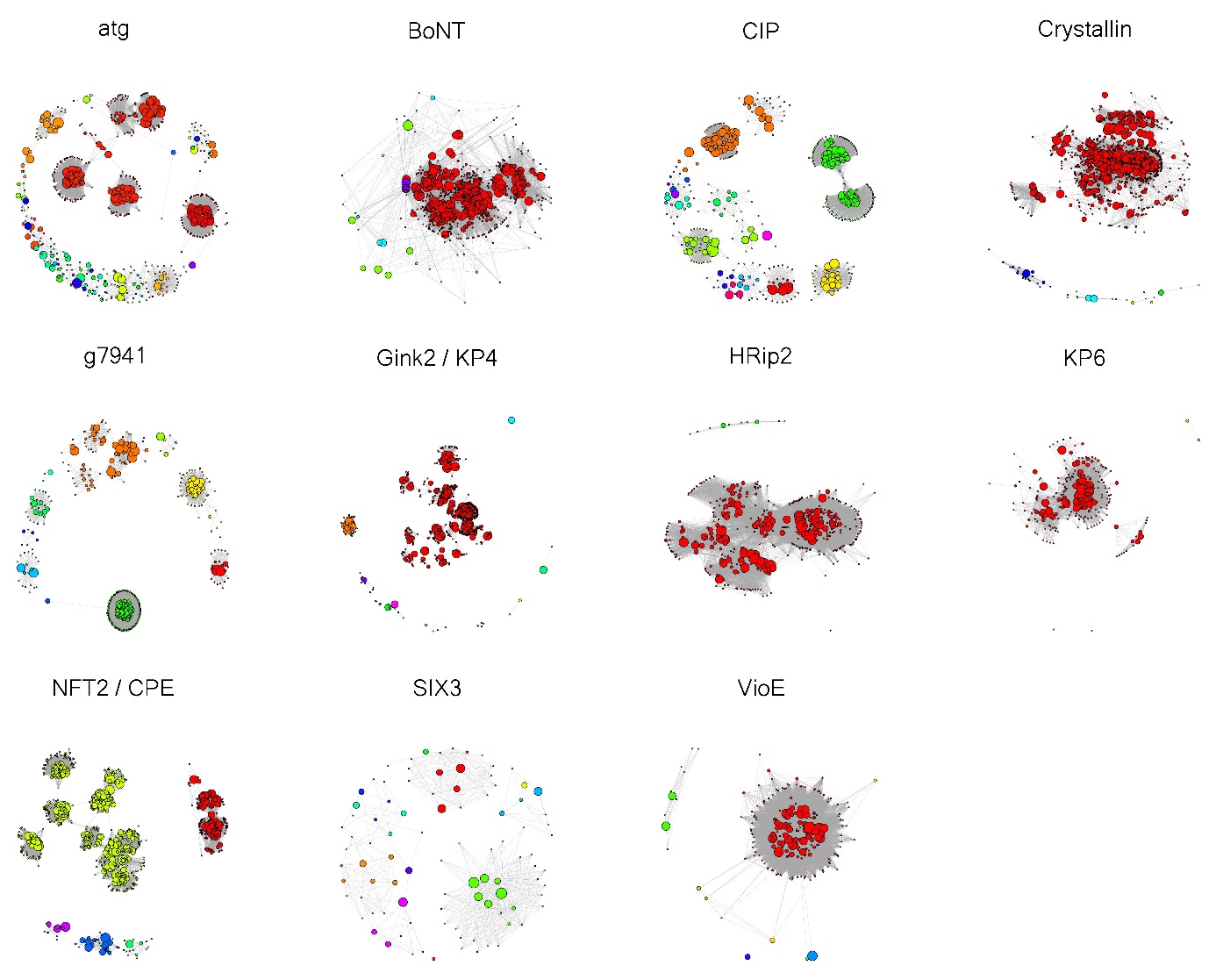

**Figure S8.** Each circle represents a different MMseqs cluster and each super-cluster is represented with a different colour. Super-clusters were identified using all v all HMM comparisons between MMseqs clusters. Nodes are connected where there is a significant HMM match between them.

We defined super-clusters of potentially evolutionarily related sequences as groups of HMMs / singletons where each member was homologous to at least one other member in the group. We found that homologues of eight of the 12 families grouped into a major ancestral super-cluster that comprised at least 60 % of the HMMs and singletons (**Fig. S9**). For instance, the largest super-cluster was found for the Crystallin fold, and it comprised 1,187 (96 %) HMMs and singletons. The second largest Crystallin fold HMM super-cluster only contained 19 HMMs and singletons. For homologues of the AltA1, BoNT, Gink2 / KP4, HRip2, KP6 and NFT2 / CPE OCE families, the largest super-clusters comprised 89 %, 94 %, 91 %, 97 %, 98 %, 69 %, and 89 %, respectively, of HMMs and singletons. HMMs and singletons comprising homologues of OCEs with atg, CIP, g7941, SIX3 and VioE folds were spread more evenly across super-clusters, with the largest super-clusters comprising from 35 % (SIX3) to 47 % (g7941) of HMMs and singletons.

An analysis focused specifically on 11,251 AltA1 family homologues showed that they formed 7 super-clusters with possible common ancestry. The largest of these clusters, super-cluster 1, was present in 10 divergent fungal classes spanning both Basidiomycetes and Ascomycetes, and it contained 627 related HMMs / singletons spanning 5,135 proteins. This super-cluster contained the PDB entries for the structurally similar proteins TSP1, AltA1 and PevD1. Other super-clusters were smaller and ranged from being broadly taxonomically distributed to highly taxonomically restricted. For instance, super-cluster 2, which contained 44 HMMs / singletons and 220 proteins, spanned four Ascomycete and one Basidiomycete class, whereas super-cluster 6 contained one HMM and one singleton, together comprising 28 proteins, and was only present in the Leotiomycetes.

In the analysis focused on AltA1, we also included the homologues of the top five PDB hits of all OCEs, which included many metazoan and bacterial proteins. Though there was structural similarity between OCEs and metazoan and bacterial proteins, notably avidin and streptavidin, we did not detect any evidence of distant common ancestry (e < 1e-5).

Community structure detection uncovered evidence for several distinct groups within AltA1 super-cluster 1. These were groups that were connected to each other by relatively few of their members, and thus common ancestry for these groups was less clear. The HMMs containing TSP1, AltA1 and PevD1 all appeared in the same sub-cluster (sub-cluster 7) of super-cluster 1, which had members in the five Ascomycete classes. This suggests that these proteins share a common ancestor and supports previous findings for AltA1 and PevD1.

Overall, HMM comparisons suggest that the majority of effector folds share distant common ancestry. However, we cannot rule out convergent *de novo* emergence of similar effector folds, or fold recruitment from unrelated sequences, especially since several of the families formed distinct clusters with no detectable homology based on HMM-HMM comparisons.

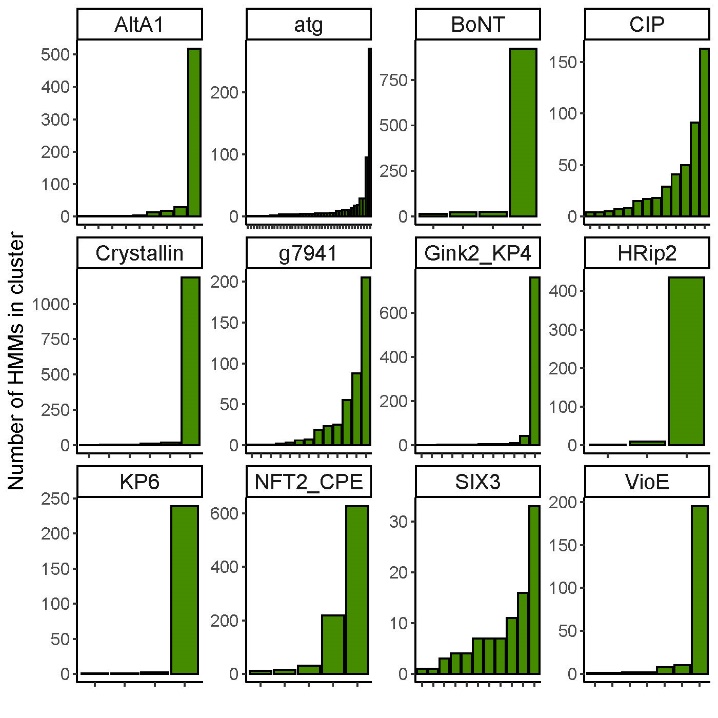

**Figure S9** All super-clusters for each family are laid out on the x axis from smallest to largest. The y axis shows the number of MMseqs clusters that make up the super-cluster.

5. **Distribution of frustrated residues in OCE proteins**

Protein structures are subjected to two contradictory evolutionary forces. On one hand, they are under pressure to maintain a thermodynamically favourable structure that folds efficiently. On the other hand, they are under pressure to maintain or adapt their functions. This evolutionary dynamic leads to a situation in which most evolved proteins fold into a stable structure where most amino acid contacts are thermodynamically optimal but a few are thermodynamically unfavourable and described as ‘frustrated’. Frustrated contacts are usually important for a protein’s function as they allow for changes in local conformation upon substrate binding.

To test whether OCEs exhibit a frustrated and evolvable surface surrounding a thermodynamically stable core, we assessed the relationship between frustration index, where a higher frustration index indicates lower levels of thermodynamic frustration for a given amino acid, and relative surface exposure (Main text **Figure 4A**). We found that across the three structural groups we tested (AltA1, KP6 and Gink2 / KP4), there was a negative correlation between relative surface exposure of amino acids and frustration index (Spearman’s rho = -0.4, Figure M1 B); there were also correlations within individual groups of -0.4 (AltA1 and KP6) and -0.31 (Gink2 / KP4). Amino acid contacts with a frustration index of less than -1 are generally considered highly frustrated, i.e. most other pairings of amino acids in this position would be more thermodynamically favourable.

We hypothesised that frustrated surface exposed residues might be in more dynamic parts of the proteins. However, the correlation between frustration and local deviation in structure was -0.17, (**Fig. S10**). Within families correlations were -0.13, -0.1 and -0.27 for AltA1, Gink2 / KP4 and KP6, respectively. There was a stronger correlation overall between relative surface exposure and local structure deviation at 0.36. This correlation was particularly high in the KP6 family at 0.63, intermediate in GNK2 / KP4 at 0.5 and lower in AltA1 at 0.3. Similar to the correlative analysis, there was a statistically weaker increase in local structural deviation among highly frustrated amino acids (P = 0.0008, **Fig. S10**).

These findings suggest that frustrated residues are distributed throughout the surfaces of OCEs but that they are not necessarily in the most structurally diverse parts of the surface. Overall, the stable core of the protein is both structurally conserved and minimally frustrated.

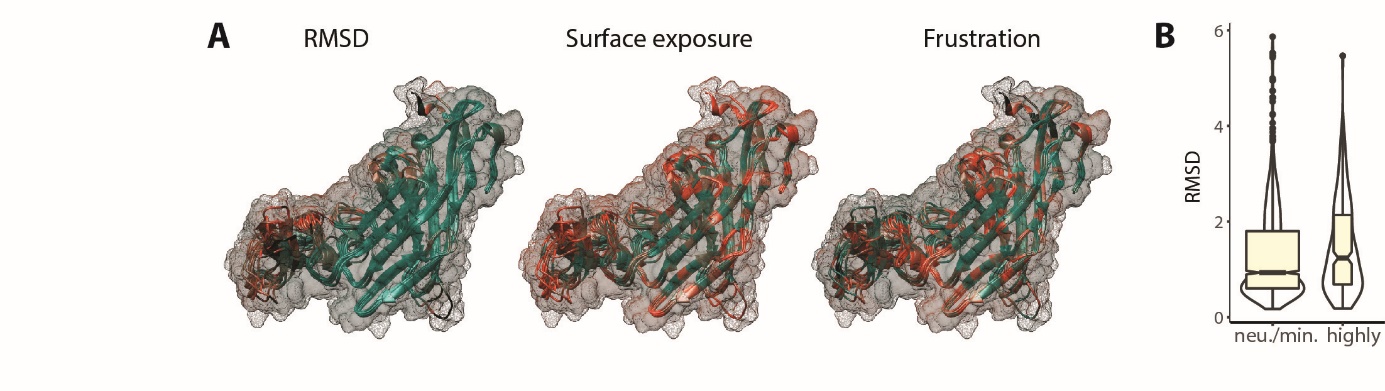

**Figure S10. Relationship between frustration and structural variation estimated using RMSD.**

To test whether amino-acids at the surface vary in frustration during evolution more than buried residues, we aligned each descendant with their respective N0 ancestor and calculated absolute delta frustration index for each residue. We then tested the correlation between average atomic distance from the surface in Angstroms and absolute change in frustration. For this, distance to protein surface was calculated using the MSMS 2.6.1 software from the ssbio python package [6]. Although there was a weak negative correlation between distance from surface and delta frustration index (Spearman’s rho = 0.2, P = 0), there was a much greater variability in delta frustration index among amino acids at the surface of the protein compared with in the core. **Figure S11** shows a quantile regression of delta frustration index on distance from the surface, 0.5, 0.75, 0.9 and 0.99 quantile regression lines are shown. There was also an overall increase in delta frustration per residue among surface exposed residues relative to buried residues (surface exposed were defined as < 2.5 angstroms distance from the surface on average – this was based on the fact that a carbon atom is about 2 angstroms wide) (**Figure S11**). Overall, OCE residues close to the surface have changed frustration patterns more during evolution than residues that are buried.

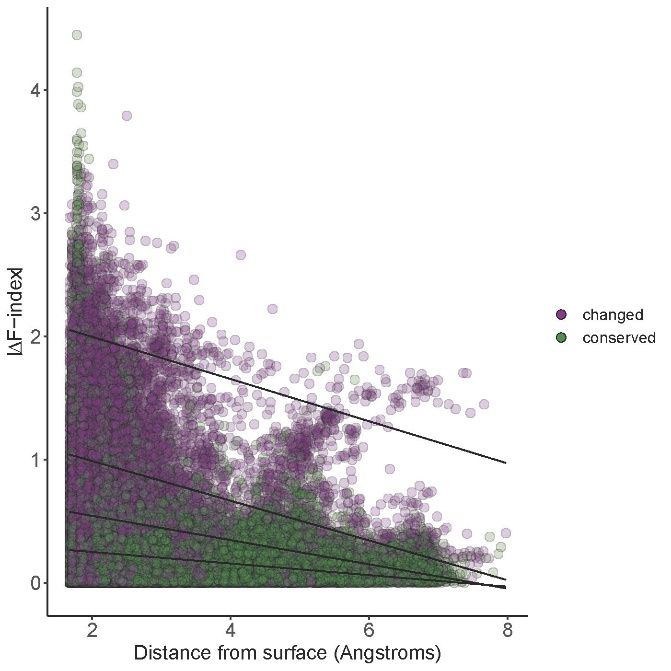

**Figure S11. Relationship between residues frustration variation (y-axis) and surface exposure (x-axis) in three OCE families.** Conserved residues are shown in green, variants in purple. 0.5, 0.75, 0.9 and 0.99 quantile regression lines are shown in black.

6. **Correlation between change in frustration relative to common ancestor and structural similarity to common ancestor**

To study the relationship between change in frustration and change in structure in OCEs, we compared all OCEs against their common ancestor at the root of the tree and subtracted Z scores from the ancestor self-match Z score to estimate structural deviation. Next, we calculated the average absolute change in frustration for any given site relative to the ancestor, and calculated sequence similarity between ancestors and their descendant OCEs. There was a significant correlation between absolute change in structure and absolute change in frustration (on average per residue) (rho = 0.59, P = 0, **Fig. S12A**) and a stronger anticorrelation between sequence similarity to the ancestor and absolute change in frustration (rho = 0.87, P = 0) (**Fig. S12B**). This indicates that mutation is a major driver of frustration change in OCEs and that frustration variation (increase and decrease in frustration altogether) has a strong impact on structure of OCEs. The distribution of frustration and structure variation along phylogenetic trees indicated that these parameters fluctuated over the evolution of OCEs (**Fig. S13**).

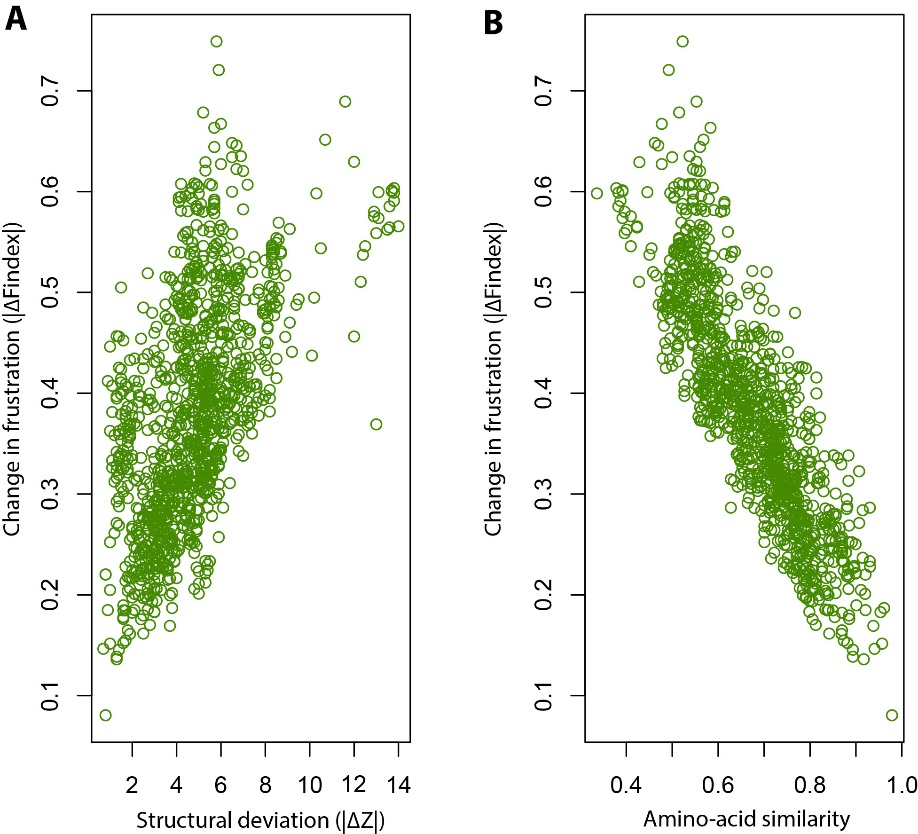

**Figure S12. Relationship between absolute change in frustration for any given site relative to the ancestor (Y-axis) and structural deviation from common ancestor (A) and calculated sequence similarity between ancestors and their descendant OCEs (B).**

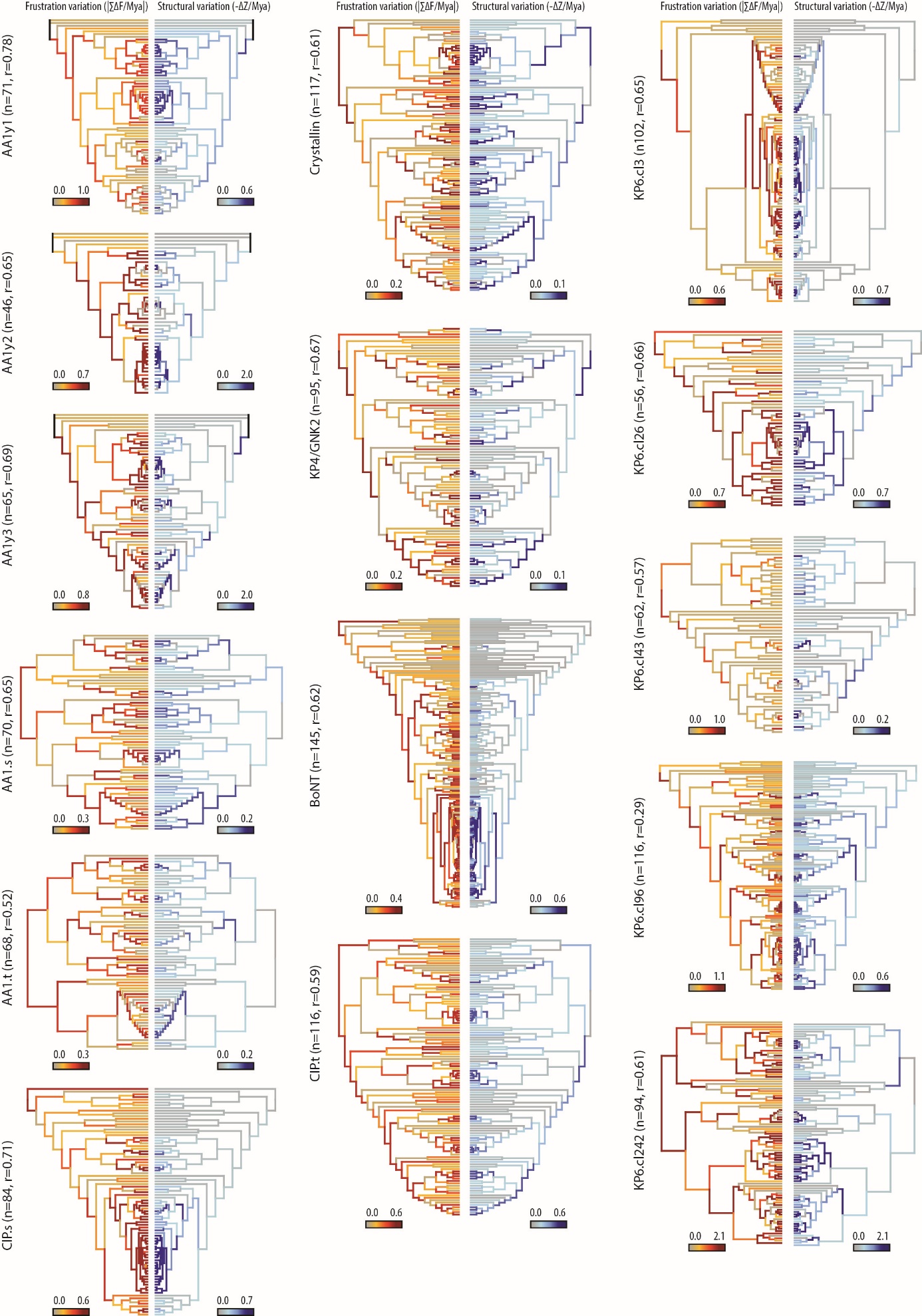

**Figure S13. Frustration (red, left side) and structural (blue, right side) variation during the evolution of 15 OCE clusters from 6 families mapped on time-calibrated phylogenies.** N is the number of modern OCEs per group, r is Pearson's product-moment correlation between frustration and structural variation across all branches of the tree. Mya, Million years ago.
